# Genomic Alterations and Possible Druggable Mutations in Carcinoma of Unknown Primary (CUP)

**DOI:** 10.1101/2021.03.25.436921

**Authors:** Hamidreza Aboulkheyr Es, Hamid Mahdizadeh, Amir Abbas Hedayati Asl, Mehdi Totonchi

## Abstract

Carcinoma of Unknown Primary (CUP) is a heterogeneous and metastatic disease where the primary site of origin is undetectable. Currently, chemotherapy is an only state-of-art treatment option for the CUP patient. Employing molecular profiling of the tumour, particularly mutation detection, offers a new treatment for CUP in a personalized fashion. Here, we analyzed mutation and copy number alterations profile of 1,709 CUP samples deposited in GENIE cohort and explored potential druggable mutations. We identified 52 significant mutated genes (SMG) among CUP samples, of which 13 (25%) of SMG were potentially targetable with drugs reproved for the know primary tumour or undergoing clinical trials. The most variants detected were *TP53* (43%), *KRAS* (19.90%), *KMT2D* (12.60%), and *CDKN2A* (10.30%). Additionally, the presence of similar variants of *TERT* promoter in CUP compared to NSCLC samples suggests these mutations may serve as a diagnostic marker for identifying the primary tumour in CUP. Taken together, analyzing mutation profiling of the CUP tumours may open a new way of identifying druggable targets and consequently administrating appropriate treatment in a personalized manner.

## Introduction

From all patient diagnosed with cancer, 3-5% present as metastatic carcinoma of unknown primary site (CUP) [1]. It is classified as any metastatic epithelial tumour where, following extensive clinical history, physical examination, radiological studies and histopathological investigations failed to identify the primary site of tumours [2]. Clinically, there has been no consensus and standard treatment guidelines that result in various chemotherapy regimens. Because of CUP tumour heterogeneity, the current clinical trials are difficult to perform, resulting in a poor prognosis with a median survival of less than 12 months and 5-year survival of 14% [3]. Thus, there is an urgent need to improve treatment modalities and prolong patients’ survival with CUP [4].

Personalized cancer medicine using genomics technologies opened new ways to treat various types of cancers using the identification of targetable mutations [5–9]. Recent studies have highlighted the crucial role of precision medicine in patient stratification and the selection of effective treatment in malignant types of cancer [10, 11]. Moreover, several studies have demonstrated improved overall survival in patients with advanced and metastatic cancers who have received genetically-matched targeted therapies [12, 13]. In CUP tumours, the implementation of this approach may improve treatment by targeting tumour-specific and druggable somatic variants in a personalized manner [3]. The AACR Project Genomics Evidence Neoplasia Information Exchange (GENIE) has recently collected the genomic information including mutations and copy number variation of the wide range of solid tumours including CUP from both primary and metastatic tumours [14–16]. Using these public data, we analyzed the genomic mutations and copy number alterations of 1,709 CUP samples to provide insight into the genetic makeup of these tumours and determined potentially druggable targets.

## Material and Methods

### Data collection

GENIE v5.0 provided the mutation, copy number variation, gene fusion and clinical information of 59,442 tumour samples [15]. Most onco-types were classified into 17 categories according to Oncotree (http://oncotree.mskcc.org/oncotree/). The onco-types not included in these 17 categories were excluded from our analysis. Raw data were downloaded from Synapse (syn17112456, https://www.synapse.org/) and provided by the GENIE project using either R commands or cbioportal (https://genie.cbioportal.org/) [17, 18]. The preprocessing protocols for these data are described in the GENIE-provided data guide.

### Significantly mutated genes (SMG) analysis

The SMG analysis performed according to the previously developed criteria and protocols 20,21. We used the MuSiC suite [19] to identify significant genes for CUP samples and also for Pan-Cancer tumours according. This test assigns mutations to seven categories: AT transition, AT transversion, CG transition, CG transversion, CpG transition, CpG transversion and indel, and then uses statistical methods based on convolution, the hypergeometric distribution (Fisher’s test P-value < 0.05), and likelihood to combine the category-specific binomials to obtain overall P-values. Notably, the genes with a cohort level alteration frequency of ⩾ 5% or a tumour type-specific alteration frequency of ⩾ 30% were included in our analysis, while tumours having no mutation, or more than 500 mutations were excluded in this study. Differentially mutated sites were plotted using Mutation-Mapper module in cBioportal. (http://www.cbioportal.org/mutation_mapper.jsp).

### Copy number variation analysis

Copy number alteration data were available at AACR Project GENIE, in cbioportal. In the present study, we selected the 17 most common cancer types for comparing their copy number variation frequencies with CUP samples. We calculated the changes in the average frequency of copy number variation (amplification and deletion) of CUP and Pan-cancer samples using provided R code in cbioportal.

### Mutual exclusivity and co-occurrence analysis

We used Fisher’s exact test to identify pairs of SMGs with significant (P-value < 0.001 by Benjamini–Hochberg) exclusivity and co-occurrence. We identified significant pairs by analyzing all CUP samples together. Then we used a de novo driver exclusivity algorithm known as Dendrix [20] to identify sets of approximately mutually exclusive mutations on all samples together. The plotting for mutual exclusivity and co-occurrence was performed using Gitools software (version 2.3.1) [21].

### Data Availability

The genomic data from the GENIE dataset used in this study are openly available for download in https://www.synapse.org, reference number [syn17112456] [15].

## Results

### Clinical characteristic of samples

In total, 45,048 samples across 17 cancer types, including CUP were included in this study. The sample type distribution was 24,567 primary and 15,484 metastasis tumours in GENIE cohorts. The hotspot regional mutations and copy number variations of these samples were available from GENIE and cBioportal. According to the information provided by GENIE, we divided samples into 17 broader cancer types, including CUP samples (Fig. 1A). The cancer categories containing the most samples were non-small cell lung cancer (9,085 (15.3%)), breast invasive ductal carcinoma (8,712 (14.7%)), colorectal cancer (5,961 (10.0%)), Glioma (3,214 (5.4%)), Melanoma (2,492 (4.2%)), prostate cancer (2,214 (3.7%)). The number of CUP samples registered in this cohort was 1709 (2.9%), dividing to 1222 metastatic (71.5%), 288 primaries (16.9%), 182 (10.6%) not applicable or heme and 17 (1.0%) unspecified (Fig. 1B). For gender information among CUP patients, 50.5 % of patients were female and 49.5% were male (Fig. 1C).

**Figure 1:**
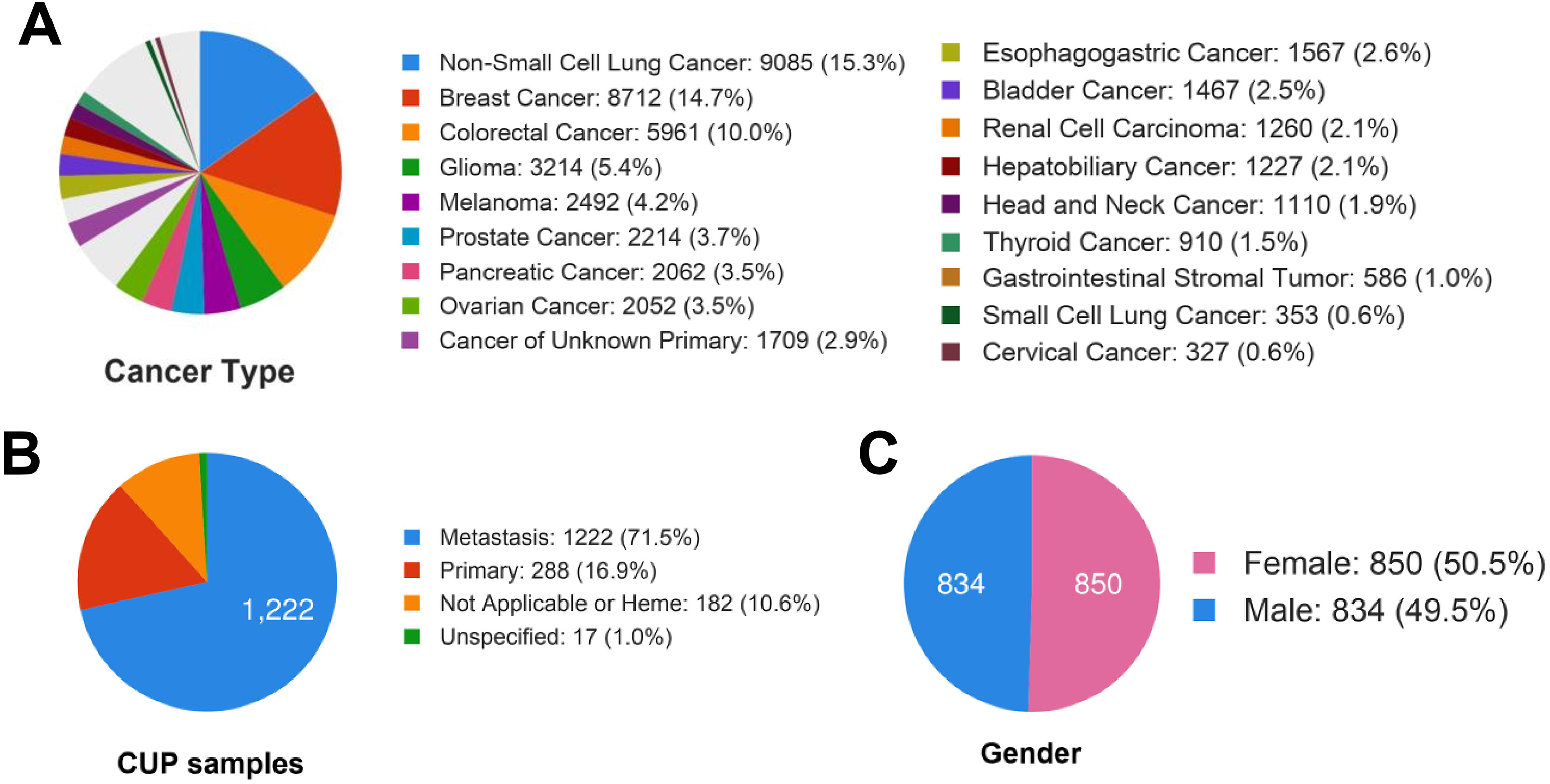
Overview of the GENIE database. Distribution of tumour types among cases successfully sequenced and analyzed in this cohort.

### Significantly mutated genes (SMG) in CUP samples

We analyzed the most genomic mutations of hotspot regions at the gene level in CUP samples in GENIE according to the previously developed method [8, 9]. In total, 52 SMG was identified (Fig. 2A; Supplementary Table 1). Among SMGs, the mutation rate of *TP53*, *KRAS*, *ARID1A*, *SMARCA4* and *KMT2D* were recorded significantly higher than other identified SMGs (Fig. 2B, Supplementary Table 1). The pathway enrichment analysis of identified SMGs resulted to the involvement of SMGs in a wide range of cellular processes, (Fig. 2C, Supplementary Table 2), including transcription factors/regulators, receptor tyrosine kinase signalling, cell cycle, IGF pathway-protein kinase B signalling, phosphatidylinositol-3-OH kinase (PI(3)K) signalling, Wnt/β-catenin signalling, *PDGF*, *FGF*, *EGF*, *TGF*-*beta*, and Notch signalling pathways and integrin signalling pathway. The identification of *MAPK, PI(3)K* and Wnt/β-catenin signalling pathways is consistent with classical cancer studies. Notably, almost all samples had at least one non-synonymous mutation in at least one SMG. The average number of point mutations in these genes varies across samples, with the highest (512 mutations for TP53 across 727 cases) and the lowest (15 mutations for *GLI3* across 15 cases) (Fig. 2B. Supplementary table 1). This suggests that the numbers of both cancer-related genes (52 identified in this study) and cooperating driver mutations required during oncogenesis are few, although large-scale structural rearrangements were not included in this analysis. Interestingly, in line with the previous study performed by Zehir et al. 2017 [8] highlighting *TERT* promoter mutations across few primary tumours, we observed similar mutation of *TERT* promoter among CUP samples (n=91) (Fig. 2D). Although the clinical relevance of mutations in the *TERT* promoter remains incompletely understood, our results reaffirm the high prevalence of these alterations in patients with advanced solid tumours and suggest an association with disease progression and poor outcome. Additionally, the presence of similar mutation of TERT promoter in CUP and NSCLC samples suggests these mutations may serve as a diagnostic marker for identification of primary tumour in CUP patients.

**Figure 2:**
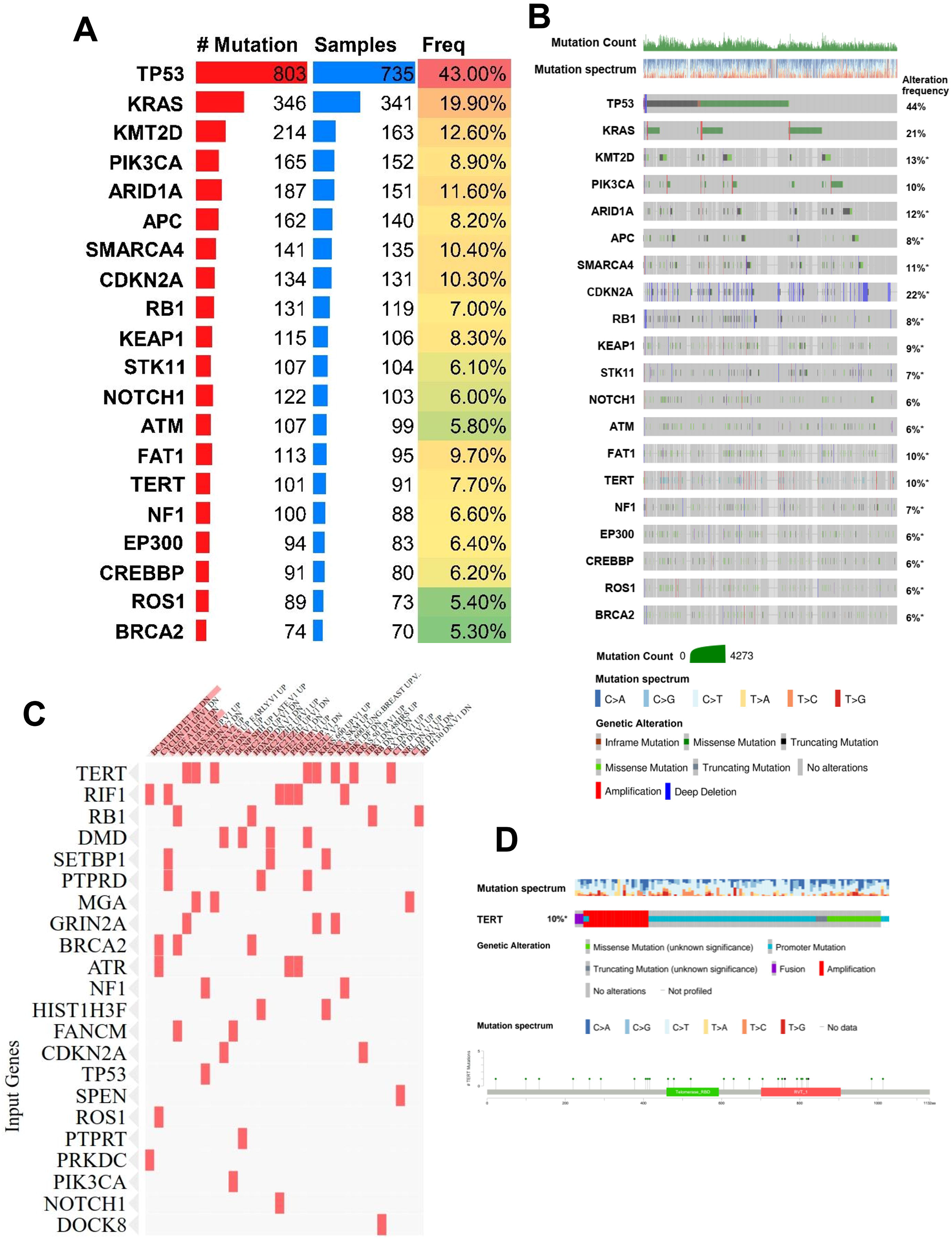
The most significant mutated genes in CUP samples. **(A)** Mutation frequency of SMGs. Genes with a cohort-level alteration frequency of > 5% or a tumour type– specific alteration frequency of > 30% are displayed. **(B)** Genomic alterations of 52 SMGs within CUP samples. **(C)** Pathway enrichment analysis of SMGs from MSigDB. **(D)** Genomic alterations identified in TERT promoter among CUP samples.

### Mutual exclusivity and co-occurrence among SMGs

The 1,035 pair-wise exclusivity and co-occurrence analysis of the 52 SMGs found 198 mutually exclusive (P-value < 0.001) and 837 co-occurring (P-value < 0.001) pairs (Fig. 3 and supplementary Table 3) among cup samples. Pairs with significant exclusivity were include *KRAS* and *FAT1*, *KRAS* and *NOTCH3*, *KRAS* and *NF1*, *KRAS* and *DMD* and *CDKN2A* and RB1 in CUP samples. Additionally, the cohort analysis identified pairs with significant co-occurrence, including KRAS and APC, TP53 and APC, KRAS and CDKN2A, KRAS and STK11, KRAS *KEAP1*, and *SMARCA4* and *KEAP1* highlighting the importance of this oncogene in cup tumours.

**Figure 3:**
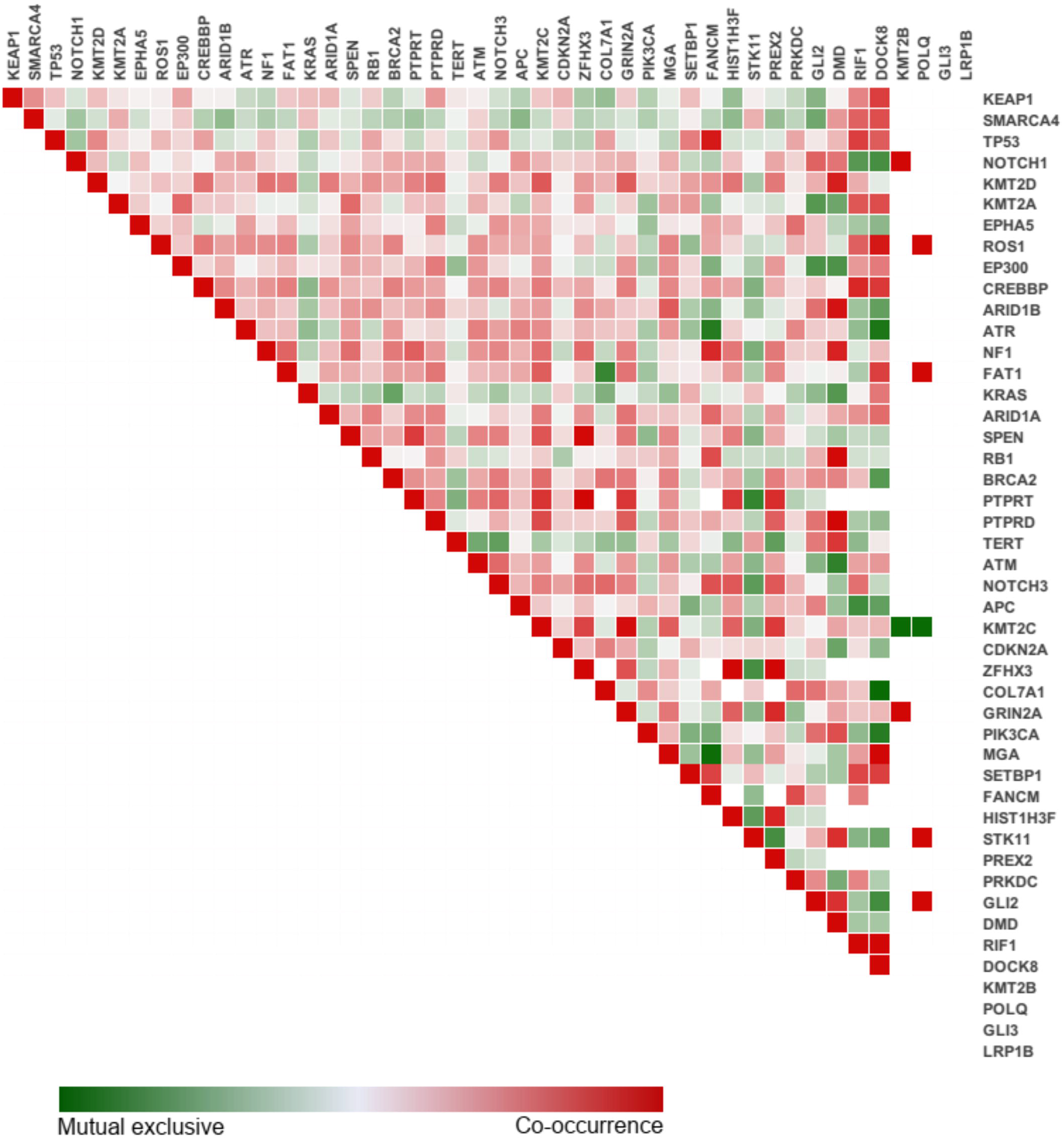
Mutual exclusivity and co-occurrence between identified SMG in CUP.

### Copy number alteration among cup samples

The copy number variation differences within cup samples resulted into identification of 624 frequently amplified/deleted regions. Significant amplification of *MYC*, *FGF4* and *FGF19* observed in a small fraction of patients (Fig. 4A) while deletion of cell cycle-related genes *CDKN2B* and *CDKN2A* were detected in only 10 and 20 percent of patients respectively (Fig. 4A). Further, we analyzed copy number alteration of the CUP-SMGs within CUP samples (Fig. 4B) and across primary tumours of 14 cancer types registered in GENIE (Fig. 4C, Supplementary Table 4). Among CUP samples, a deep deletion of *TP53*, *RB1*, *CDKN2A*, and *STK11and* amplification of *KRAS* and PIK3CA were observed. In a pan-cancer analysis, amplification of *KRAS* and *PIK3CA* in the breast (66 and 114 of cases) and non-small cell lung cancer (46 and 48 of cases), *TERT* in non-small cell lung cancer (114 of cases) and *ATR* in breast cancer (36 of cases), were the most amplified genes, while deletion in *CDKN2A* in glioma (676 of cases), *RB1* and *TP53* in small cell lung cancer (15 of cases) were observed in these 14 different cancer types (Fig. 4C). Among these genes with significantly altered copy numbers between CUP and primary tumours, a significant amplification of *TERT* promoter was observed in both CUP and non-small cell lung cancer samples compared to glioma and breast primary tumours suggesting that copy number variation of *TERT* may play diagnostic role for identification of origin of CUP tumours (Fig. 4D).

**Figure 4:**
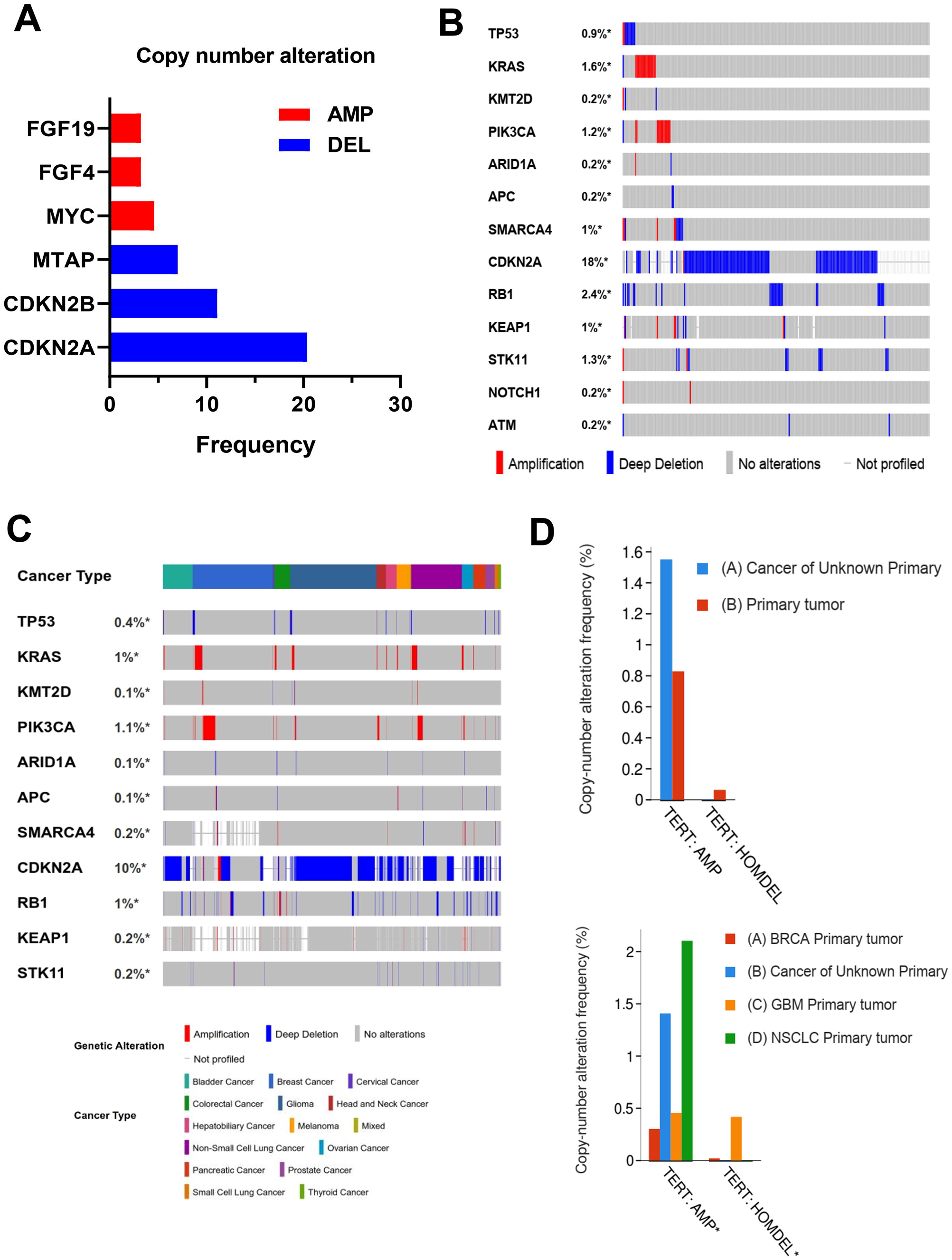
Copy number variation among CUP and other cancer types. **(A)** major copy number alteration detected in CUP samples. **(B)** Amplification and deletion status of identified CUP-SMGs within CUP samples and **(C)** other known primary tumours registered in GENIE database. **(D)** copy number variation analysis of TERT between CUP and all primary tumours (up panel) and NSCLC, GBM and BRCA (bottom panel).

### Mutation frequency of CUP-SMGs across 17 known primary tumours

To identify similar and targetable mutation pattern in CUP, we analyzed and compared genomic alteration frequency of identified CUP-SMGs in primary tumour types across 17 cancer types in GENIE (Fig. 5A). The majority of CUP-SMGs mutations were enriched in non-small cell lung cancer (4,221 cases) colon cancer (4,011 cases) and breast cancer (3,376 cases) (Fig. 5A).

**Figure 5:**
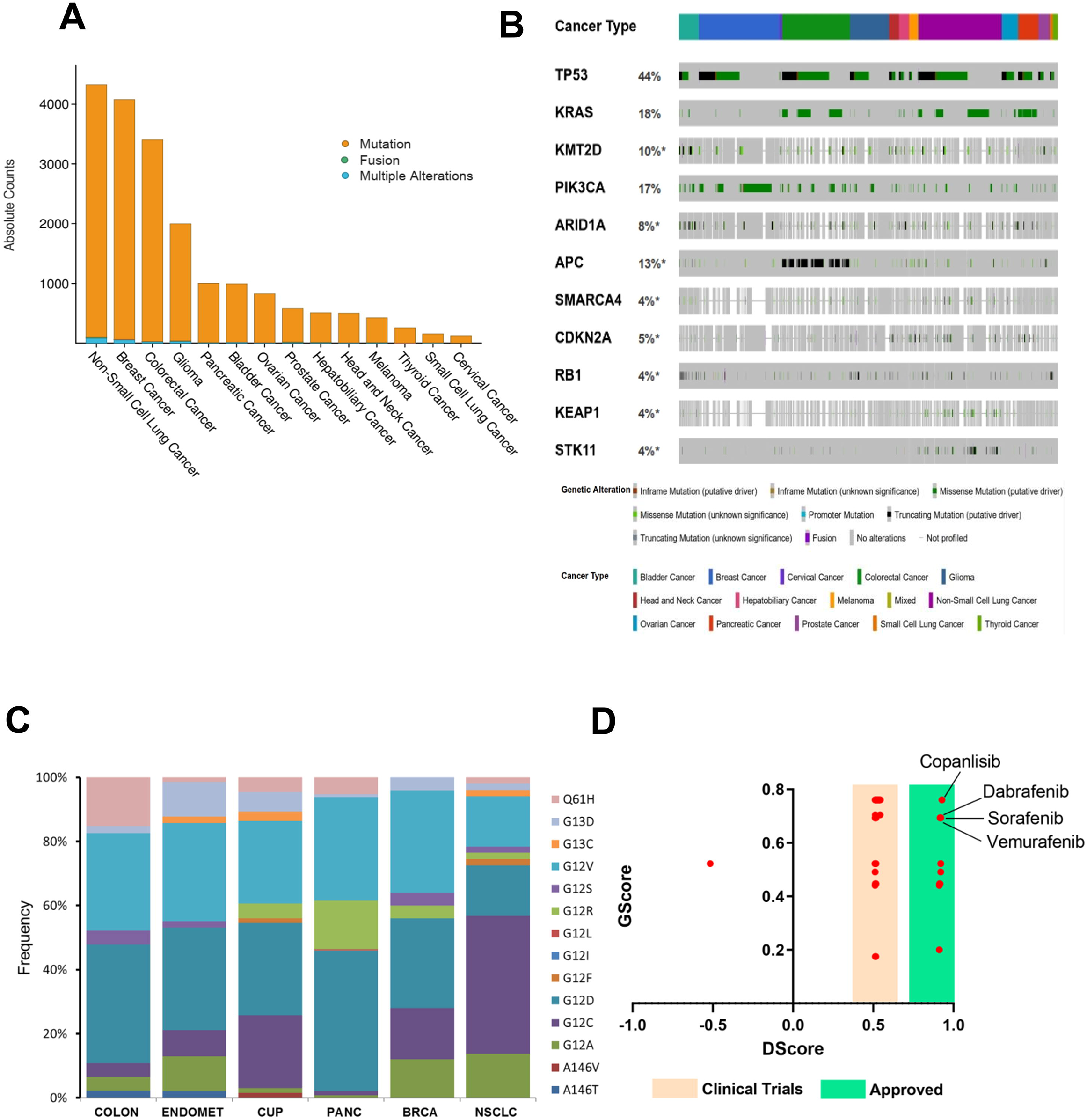
Mutation frequency of CUP-SMGs across 17 cancer types. **(A, B)** Distribution of genomic alterations in 52 CUP-SMGs across primary tumours of 17 different cancer types in GENIE cohort. **(C)** Distribution of hotspot mutations identified in *KRAS* among six different cancer types including CUP. **(D)** The results of gene-drug association analysis using PanDrug platforms. The best candidate drugs with highest GScore and Dscore are labeled.

The most frequently mutated gene in this cohort was TP53 (44% of total samples) (Fig. 5B). Its mutations predominate in non-small cell lung cancer (46.36%; 2,517 cases), colon cancer (65.55%; 2,365 cases) and breast cancer (36.26%; 2,060 cases) (Fig. 5B). *KRAS* is the second most commonly mutated genes, occurring frequently (>10%) in most cancer types (pancreatic:74.6%, colon cancer:44.24%, non-small cell lung cancer:30.93%) except hepatobiliary carcinoma, cervical cancer, bladder cancer, thyroid cancer, melanoma, small-cell lung cancer, head and neck carcinoma, prostate and breast cancer (Fig. 5B). *PIK3CA* mutations was frequented in breast cancer (36.7%) and cervical cancer (25.14%), being specifically enriched in luminal subtype tumours. Many cancer types carried mutations in chromatin re-modelling genes. In particular, histone-lysine N-methyltransferase genes *KMT2D*, *KMT2C* and *KMT2B* in bladder, lung and endometrial cancers, whereas the *KMT2A* is mostly mutated in non-small cell lung cancer and colon cancer. Mutations in *ARID1A* are frequent in non-small cell lung cancer, colon cancer, bladder cancer and breast cancer, whereas mutations in *KEAP1* and *STK11* was predominate in non-small cell lung cancer (8.62% and 11.75% respectively) (Fig. 5B). *KRAS* mutations are typically mutually exclusive, with recurrent activating mutations (*KRAS* (Gly 12) and *KRAS* (Gly 13) common in colon cancer, non-small cell lung cancer and pancreatic cancer. We compared the most common hotspot mutations in *KRAS* between CUP and other *KRAS* mutation enriched cancer types (Fig. 5C). Comparing hotspot mutations resulted to enrichment of G12D and G12R in pancreatic cancer, G12C, G12F and G13C in non-small cell lung cancer and CUP samples. These data highlight similarity of KRAS hotspot mutations between CUP and NSCLC.

### Targetable mutations and drug candidates

To identify or predict possible therapeutics based on genomic alterations identified in SMG in CUP samples, we performed a gene-drug association analysis using PanDrugs platforms [22]. The gene-drug associations classified into two groups called “Drug targets” in which drugs can directly target genes that contribute in disease phenotype, and “Biomarkers” where genes are representing a drug-response associated status while its protein products are not targetable [22]. From 262 identified interactions, 8.7 % (23/262) was classified as a direct drug target, while 91.3 % (239/262) of gene-drug interactions identified as Biomarker (Fig. 5D). Interestingly, we found five FDA approved drugs, Crizotinib (GScore: 0.76. Dscore: 0.95) and Copanlisib (GScore: 0.76. Dscore: 0.92), Debrafenib, Sorafenib, Vemurafenib, and Regorafenib as best candidate for targeting ALK/MET, PIK3CA, and BRAF inhibitors respectively (Fig. 5D. Supplementary Table 5). Moreover, various off-label and clinically investigating compounds for targeting mutated KRAS were identified, although the GScore and DScore of these compounds did not reach a high score (Supplementary Table-5). Everolimus (mTOR inhibitor), Bortezomib (26S proteasome inhibitor), and Pemetrexed (chemotherapy agent), were identified with the highest GScore and DScore compared to the other drugs candidates in this group (Fig. 5D. Supplementary Table-5). Taken together, these data highlight presence of at least one druggable variants and potential of using genomic alteration guided targeted therapy in CUP patients.

## Discussion

Currently, combination chemotherapy regimens have been considered as first-line of therapy for CUP patients [23]. Personalized cancer therapy using the identification of druggable mutations has encouraged mutational profiling of various types of tumours including metastasis tumours, including CUP [24–26]. This study analyzed the most significant mutated genes and identified the most prevalence variants in 1709 CUP samples. The gene-drug association studies suggested that at least one of the identified variants is linked to the known oncogenic driver mutations and approved targeted therapy or therapeutics are currently in clinical trial studies highlighting the potential of genomic alteration-based treatment approach for a patient with CUP. In line with this concept, numerous clinical studies have been reported durable treatment responses using mutation-matched targeted therapies drugs, including EGFR, BRAF, KIT, and MET [12, 27–29].

Currently, targeted therapy agents Crizotinib and Copanlisib approved for the treatment of tumours that harbour mutations in ROS1/MET/ALK and PIK3CA, while therapeutic agents for the other identified variants including FGFR family, MYC, MET, and KRAS are currently being investigated in active and ongoing clinical trials. A large proportion of the mutations detected in this study are known to be associated with various signal transduction pathways, apoptotic regulation, cell cycle progression, and receptor tyrosine kinase signalling regulations. These results can be promising because the majority of available targeted drugs act through targeting one of these pathways, which are commonly altered in various types of known tumours [30–34]. The most mutated gene identified in this study was TP53 (43%, 743/1709) with numerous different non-synonymous coding region variants. Similar to these data, previous studies demonstrated the association of TP53 mutations in metastatic progression in multiple cancer types, supporting the presence of high mutation load on TP53 reported in CUP [35, 36].

Other common variants detected in this cohort were observed in genes involved in the activation and regulation of key signal transduction pathways, including BRAF and KRAS. This is the first study to report various codon 12 variants of KRAS in CUP samples. The detection of codon 12 mutations in this cohort is consistent with the highly aggressive behaviours of CUP tumours [24, 28]. Furthermore, characterizing the mutational status of KRAS has become clinically relevant in some malignancies, because the presence of a KRAS mutation is known to stimulate resistance to some tyrosine kinase inhibitors [37–40]. Although, currently no approved therapeutic agent to target and inhibit mutant KRAS activity available, however, recent clinical studies reported a partial response in CUP patients with a KRAS(G12D) mutation treated with trametinib (MEK inhibitor) [29, 41]. In this study, we also observed KRAS(G12C) variant in 25% of CUP samples. Recent promising results from Sotorasib (AMG-510); a specific covalent inhibitor of K-RAS(G12C) in NSCLC suggest the detection of these variants of KRAS as a possible druggable target in CUP patients [42]. Moreover, targeting KRAS(G12C) using Sotorasib in advanced solid tumours showed an encouraging anticancer activity which might be useful in CUP [43]

Similar to other studies, we also identified an activating BRAF V600E mutations in 4.3% (74/1709) cases [23–25]. This offers the potential of using BRAF inhibitors such as Vemurafenib and Dabrafenib for CUP with BRAF (V600E) mutations. In line with these, through the gene-drug association analysis, we also observed a high GScore and DScore of BRAF inhibitors Dabrafenib and Vemurafenib for targeting V600E variant identified in CUP samples. Moreover, a clinical study showed a complete clinical response of CUP patients treated with BRAF(V600E) targeted therapy Vemurafenib in combination with immunotherapy agent Ipilimumab [44].

Mutations in MET and ERBB2 (HER2) were detected in 30 and 55 of cases respectively, suggesting the possibility of targeting these receptor tyrosine kinases [27]. Targeting MET using Crizotinib combined with HER2 inhibitor Trastuzumab has been shown with success in CUP patients. The current success of HER2 and MET targeted therapies using Trastuzumab and Crizotinib in a combination manner in advanced and metastatic tumours including HER2- and MET-mutant CUP tumour suggest the future evaluation of these genes as druggable targets in patients with CUP [45]. Our results support those of other CUP studies highlighting the value of sequencing techniques, particularly gene mutation detection, to identify actionable targets [10, 23–26].

Taken together, these data highlight the molecular heterogeneity of CUP tumours. The mutations detected across the majority of CUP cases included in this study highlights not only the genomic instability present in these tumours but also potential application of targeted therapies for a significant proportion of patients with CUP which might improve the prognosis and therapeutic decisions for the patient with CUP [11].

## Acknowledgements

The authors would like to thank GENIE for data collection, as well as cBioPortal (http://www.cbioportal.org, version v3.2.11) for the provision of data processing and customizable functions.

## Author Contributions

HAE participated in the study design and performed data extraction and analysis. He wrote the first draft of the manuscript additionally. HM participated in the study design and critically edited the manuscript. AHA participated as clinical oncologists and scientific advisor in study design, MT also participated in the study design and editing of the manuscript and he has supported the project financially.

## Availability of data and materials

All data generated and described in this article are available from the corresponding web servers and portal and are freely available to download for noncommercial purposes, without breaching participant confidentiality.

## Ethics approval and consent to participate

Not applicable.

## Consent for publication

Not applicable

## Reference

1. Greco, F.A., Cancer of unknown primary site: still an entity, a biological mystery and a metastatic model. Nat Rev Cancer, 2014. 14(1): p. 3–4.

2. Oien, K.A. and J.L. Dennis, Diagnostic work-up of carcinoma of unknown primary: from immunohistochemistry to molecular profiling. Ann Oncol, 2012. 23 Suppl 10: p. x271–7.

3. Moran, S., et al., Precision medicine based on epigenomics: the paradigm of carcinoma of unknown primary. Nat Rev Clin Oncol, 2017. 14(11): p. 682–694.

4. Hainsworth, J.D. and F.A. Greco, Cancer of Unknown Primary Site: New Treatment Paradigms in the Era of Precision Medicine. Am Soc Clin Oncol Educ Book, 2018. 38: p. 20–25.

5. Frampton, G.M., et al., Development and validation of a clinical cancer genomic profiling test based on massively parallel DNA sequencing. Nat Biotechnol, 2013. 31(11): p. 1023–31.

6. Campbell, J.D., et al., Distinct patterns of somatic genome alterations in lung adenocarcinomas and squamous cell carcinomas. Nat Genet, 2016. 48(6): p. 607–16.

7. Hoadley, K.A., et al., Cell-of-Origin Patterns Dominate the Molecular Classification of 10,000 Tumors from 33 Types of Cancer. Cell, 2018. 173(2): p. 291–304 e6.

8. Zehir, A., et al., Mutational landscape of metastatic cancer revealed from prospective clinical sequencing of 10,000 patients. Nat Med, 2017. 23(6): p. 703–713.

9. Kandoth, C., et al., Mutational landscape and significance across 12 major cancer types. Nature, 2013. 502(7471): p. 333–339.

10. Varghese, A.M., et al., Clinical and molecular characterization of patients with cancer of unknown primary in the modern era. Ann Oncol, 2017. 28(12): p. 3015–3021.

11. Rassy, E. and N. Pavlidis, Progress in refining the clinical management of cancer of unknown primary in the molecular era. Nat Rev Clin Oncol, 2020. 17(9): p. 541–554.

12. Tan, D.S., et al., Molecular profiling for druggable genetic abnormalities in carcinoma of unknown primary. J Clin Oncol, 2013. 31(14): p. e237–9.

13. Robinson, D.R., et al., Integrative clinical genomics of metastatic cancer. Nature, 2017. 548(7667): p. 297–303.

14. Micheel, C.M., et al., American Association for Cancer Research Project Genomics Evidence Neoplasia Information Exchange: From Inception to First Data Release and Beyond-Lessons Learned and Member Institutions’ Perspectives. JCO Clin Cancer Inform, 2018. 2: p. 1–14.

15. Consortium, A.P.G., AACR Project GENIE: Powering Precision Medicine through an International Consortium. Cancer Discov, 2017. 7(8): p. 818–831.

16. Litchfield, K., S. Turajlic, and C. Swanton, The GENIE Is Out of the Bottle: Landmark Cancer Genomics Dataset Released. Cancer Discov, 2017. 7(8): p. 796–798.

17. Cerami, E., et al., The cBio cancer genomics portal: an open platform for exploring multidimensional cancer genomics data. Cancer Discov, 2012. 2(5): p. 401–4.

18. Gao, J., et al., Integrative analysis of complex cancer genomics and clinical profiles using the cBioPortal. Sci Signal, 2013. 6(269): p. pl1.

19. Dees, N.D., et al., MuSiC: identifying mutational significance in cancer genomes. Genome Res, 2012. 22(8): p. 1589–98.

20. Vandin, F., E. Upfal, and B.J. Raphael, De novo discovery of mutated driver pathways in cancer. Genome Res, 2012. 22(2): p. 375–85.

21. Perez-Llamas, C. and N. Lopez-Bigas, Gitools: analysis and visualisation of genomic data using interactive heat-maps. PLoS One, 2011. 6(5): p. e19541.

22. Pineiro-Yanez, E., et al., PanDrugs: a novel method to prioritize anticancer drug treatments according to individual genomic data. Genome Med, 2018. 10(1): p. 41.

23. Gatalica, Z., et al., Comprehensive tumor profiling identifies numerous biomarkers of drug response in cancers of unknown primary site: analysis of 1806 cases. Oncotarget, 2014. 5(23): p. 12440–7.

24. Ross, J.S., et al., Comprehensive Genomic Profiling of Carcinoma of Unknown Primary Site: New Routes to Targeted Therapies. JAMA Oncol, 2015. 1(1): p. 40–49.

25. Loffler, H., et al., Molecular driver alterations and their clinical relevance in cancer of unknown primary site. Oncotarget, 2016. 7(28): p. 44322–44329.

26. Tothill, R.W., et al., Massively-parallel sequencing assists the diagnosis and guided treatment of cancers of unknown primary. J Pathol, 2013. 231(4): p. 413–23.

27. Stella, G.M., et al., MET mutations in cancers of unknown primary origin (CUPs). Hum Mutat, 2011. 32(1): p. 44–50.

28. Palma, N.A., et al., Durable Response to Crizotinib in a MET-Amplified, KRAS-Mutated Carcinoma of Unknown Primary. Case Rep Oncol, 2014. 7(2): p. 503–8.

29. Kato, S., et al., Utility of Genomic Analysis In Circulating Tumor DNA from Patients with Carcinoma of Unknown Primary. Cancer Res, 2017. 77(16): p. 4238–4246.

30. Holohan, C., et al., Cancer drug resistance: an evolving paradigm. Nat Rev Cancer, 2013. 13(10): p. 714–26.

31. Ramirez, M., et al., Diverse drug-resistance mechanisms can emerge from drug-tolerant cancer persister cells. Nat Commun, 2016. 7: p. 10690.

32. Gottesman, M.M., Mechanisms of cancer drug resistance. Annu Rev Med, 2002. 53: p. 615–27.

33. Yuan, T.L. and L.C. Cantley, PI3K pathway alterations in cancer: variations on a theme. Oncogene, 2008. 27(41): p. 5497–510.

34. Roberts, P.J. and C.J. Der, Targeting the Raf-MEK-ERK mitogen-activated protein kinase cascade for the treatment of cancer. Oncogene, 2007. 26(22): p. 3291–310.

35. Powell, E., D. Piwnica-Worms, and H. Piwnica-Worms, Contribution of p53 to metastasis. Cancer Discov, 2014. 4(4): p. 405–14.

36. Tang, Q., et al., Mutant p53 on the Path to Metastasis. Trends Cancer, 2020. 6(1): p. 62–73.

37. Pao, W., et al., KRAS mutations and primary resistance of lung adenocarcinomas to gefitinib or erlotinib. PLoS Med, 2005. 2(1): p. e17.

38. Del Re, M., et al., Contribution of KRAS mutations and c.2369C > T (p.T790M) EGFR to acquired resistance to EGFR-TKIs in EGFR mutant NSCLC: a study on circulating tumor DNA. Oncotarget, 2017. 8(8): p. 13611–13619.

39. Ohashi, K., et al., Lung cancers with acquired resistance to EGFR inhibitors occasionally harbor BRAF gene mutations but lack mutations in KRAS, NRAS, or MEK1. Proc Natl Acad Sci U S A, 2012. 109(31): p. E2127–33.

40. Misale, S., et al., Emergence of KRAS mutations and acquired resistance to anti-EGFR therapy in colorectal cancer. Nature, 2012. 486(7404): p. 532–6.

41. Ross, S.J., et al., Targeting KRAS-dependent tumors with AZD4785, a high-affinity therapeutic antisense oligonucleotide inhibitor of KRAS. Sci Transl Med, 2017. 9(394).

42. Canon, J., et al., The clinical KRAS(G12C) inhibitor AMG 510 drives anti-tumour immunity. Nature, 2019. 575(7781): p. 217–223.

43. Hong, D.S., et al., KRAS(G12C) Inhibition with Sotorasib in Advanced Solid Tumors. N Engl J Med, 2020. 383(13): p. 1207–1217.

44. Roe, O.D. and S.G. Wahl, The undifferentiated carcinoma that became a melanoma: Re-biopsy of a cancer of an unknown primary site: a case report. J Med Case Rep, 2017. 11(1): p. 82.

45. Clynick, B., et al., Genetic characterisation of molecular targets in carcinoma of unknown primary. J Transl Med, 2018. 16(1): p. 185.

